# Melatonin serves as a novel treatment in cystic fibrosis and inhibits cystic fibrosis through TGF-β1/Smad and EMT

**DOI:** 10.1101/2023.11.15.567324

**Authors:** Yang Zhang, Sun Gong, Weixin He, Jie Yuan, Di Dong, Jialong Zhang, Haomin Wang, Binghai Chen

**Author notes:** Correspondence authors: Binghai Chen (BC). These authors contributed equally to this work (Yang Zhang, Sun Gong). These authors also contributed equally to this work (Weixin He, Jie Yuan, Di Dong, Jialong Zhang, Haomin Wang).

## Abstract

**Background:** Melatonin (MEL) is an indole amine molecule primarily produced in the pineal gland. Melatonin has been shown in numerous studies to have anti-fibrosis characteristics in the kidney, liver, and other organs. However, it is still unclear how melatonin works in bladder fibrosis. We explored how melatonin affected animals with bladder fibrosis and its underlying mechanisms.

**Materials and Methods:** MEL was used to treat human bladder smooth muscle cells (HBdSMCs) after they were stimulated with TGF-β1 in vitro. Proteomic analysis and then bioinformatic analysis based on the alterations in these proteins were then performed on HBdSMCs from the different processing methods. To construct an in vivo bladder fibrosis model, we injected protamine sulfate (PS) and lipopolysaccharide (LPS) twice a week into the rat bladder for six weeks. After two weeks of PS/LPS treatment, the treatment group was treated with MEL (20mg/kg/d) for 4 weeks. Finally, we detected the expression of fibrosis markers from different perspectives. The TGF-β1/Smad pathway, and EMT in cell and bladder tissues were also identified. Further proteomic analysis was also performed.

**Results:** In the in vitro experiment, we found that TGF-β1 treatment enhanced the fibrosis markers Collagen III and α-SMA of HBdSMCs. E-cadherin expression decreased while TGF-β1/Smad pathway was activated. Vimentin and N-cadherin expressions were also elevated at the same time. Similar findings were observed in the LPS group. After MEL treatment, the expression of collagen III and α-SMA decreased, and the expression of E-cadherin increased, while the expression of Vimentin and N-cadherin also decreased. CCN1 and SQLE may be the important proteins in the development of bladder fibrosis, according to quantitative proteomics analysis. MEL can decrease their expressions which leadis to relief of bladder fibrosis. Bioinformatics analysis shows that the extracellular space structure related to metabolic pathways, actin filament binding, and stress fibers can serve as a pivotal focus in the management of fibrosis.

**Conclusion:** Melatonin attenuates bladder fibrosis by blocking the TGF-β1/Smad pathway and EMT. CCN1 appears to be a possible therapeutic target for bladder fibrosis.

## 1. Introduction

Fibrosis is a progressive disease feature with tissue scarring, which is a typical and inevitable pathological outcome of many long-lasting inflammatory conditions. Fibrosis is closely associated with bladder dysfunction including neurogenic bladder and bladder outlet obstruction (BOO) (1, 2). In BOO patients, higher pressure in the bladder causes bladder detrusor hypertrophy. Bladder morphology may change irreversibly in untreated BOO, leading to increased collagen accumulation and bladder fibrosis(3).

The pleiotropic cytokine, transforming growth factor (TGF), regulates a diverse array of biological processes, including the development of the extracellular matrix (ECM), differentiation of cells, and immunological control(4). TGF-β, the major fibrosis factor, is produced mostly by macrophages that emerge in the early stages of the wound healing response and has both anti-inflammatory and pro-fibrotic properties. In the liver, lung, kidney, skin, and heart, the development of fibrosis is correlated with the synthesis of TGF-β. The development of fibrosis has been demonstrated to be inhibited by TGF-β1 signaling pathway suppression in numerous animal models (5), and the TGF-β/Smad signaling pathway has drawn increased attention as a potent target for anti-fibrosis therapy(6).

The epithelial-mesenchymal transition (EMT) is regulated by EMT-TF, which suppress epithelial genes while activating the expression of mesenchymal components. Extracellular signals control EMT-TF expression, which frequently entails collaboration between several signaling pathways that modify epithelial plasticity in an environment-dependent manner. While a variety of signals can regulate EMT, transforming growth factor-β (TGF-β) often plays a dominant role(7–9). The EMT is recognized as a pivotal cellular mechanism contributing to the progression of tumors, fibrosis, and wound healing. Epithelial cells abandon their adhesion while adopting mesenchymal traits, including anteroposterior polarity, migratory capabilities, and the ability to invade adjacent tissues(10). Basement membranes, other extracellular matrix (ECM) structures, and cell interaction with nearby cells are all altered by these alterations.

MEL is a small indoleamine molecule (Fig S1A). It is primarily synthesized by the pineal gland during physiologically normal conditions when the hypothalamus suprachiasmatic nucleus is activated at night(11). Abundant research indicates that melatonin demonstrates anti-fibrotic effects in the liver, lungs, and various other organs (12). However, the mechanism of melatonin’s anti-fibrotic effect has remained elusive. So far, there have been no reports on the function of melatonin in bladder fibrosis. Hence, our investigation focuses on elucidating melatonin’s inhibitory impact on rat bladder fibrosis as well as its molecular mechanisms.

To validate our hypothesis, a series of experiments were conducted both in vivo and in vitro. The HBdSMCs with fibrosis phenotype stimulated by TGF-β1 were treated with different treatments of melatonin. To understand the role of melatonin in bladder fibrosis, TGF-β1/Smad pathway and EMT, we used various methods to identify and evaluate the underlying mechanism in vivo and in vitro.

## 2. Materials and methods

### 2.1 Materials

DMSO was acquired from Solarbio Technology (Beijing, China). Melatonin (purity >95%, molecular weight: 232.28) and Protamine sulfate (PS)was procured from Sangon Bioengineering (Shanghai, China). TGF-β1 was acquired from Ucallm (Wuxi, China). Lipopolysaccharide (LPS) was acquired from Beyotime Biotechnology (ST1470, Reagent grade, purified by phenol extraction, Shanghai, China). DMEM, PBS, and FBS were acquired from XP Biomed, (Shanghai, China). HBdSMCs were cultured at 37 °C, 5% CO2, and 95% air. The antibodies employed included: Phospho-Smad2 (cat. AF3362, Affinity), Phospho-Smad3 (cat. AF3449, Affinity), Smad2/3 (cat. AF6367, Affinity), E-cadherin (cat. 13116, CST), Vimentin (cat. 60330, Proteintech), Collagen Type III (cat. 22734, Proteintech), SQLE (cat. 12544, Proteintech), N-cadherin (cat. 3195, CST), α-Smooth Muscle Actin (cat. 14395, CST), TGF-β1 (cat. 21898, Proteintech), CCN1 (cat. AF3362, Affinity).

### 2.2 Animals

The Jiangsu University’s Ethics Committee and Animal Experiment Committee gave their permission to the study’s protocol. 18 female Sprague-Dawley (SD) 8-week-old rats, weighing around 200 grams, were used in the experiment. Prior to the experiments, all rats were acclimatized for one week in an environment with a temperature control of (21° C) and a light-dark cycle of 12:12 hours. Animals had freely liberty to consume enough food and water. The method described by Chae-Min Ryu et al.(13) was used to create a rat fibrosis model. All rats were randomly assigned to one of three groups. For bladder emptying and drug administration, rats were catheterized using a 1 mm epidural catheter after isoflurane anesthesia. The sham-operated group was perfused with 500 µL (sham group, n=6); the PS/LPS treatment group was first perfused with PS (10 mg/rat, dissolved in 500 µL PBS) to remove the epithelium, and the bladder was drained of PS by squeezing it after 45 min, followed by bladder rinsing with PBS. Finally, LPS (750 μg/rat, dissolved in 500 µL PBS) was instilled into the bladder through a catheter using a syringe and held for 30 min and drained by bladder pressure and rinsed with PBS after half an hour, twice a week (LPS group, n=6). The treatment group used the same PS/LPS infusion method, but after two weeks of PS/LPS infusion, a specific dose of melatonin (20 mg/kg/day dissolved in 1% ethanol) was injected intraperitoneally every day for 4 weeks (LPS + MEL group, n=6).

### 2.3 Cell Culture and Treatments

HBdSMCs were procured from Otwo Biotech Inc. (Guangzhou, China). To analyze melatonin’s impact on TGF-β1-induced HBdSMCs. First, we exposed the cells to TGF-β1 at different levels (5, 10, and 20 ng/mL) for 48 and 72 hours. Following the identification of the optimal concentration (10 ng/mL) and duration (72h) of TGF-β1, we investigated the therapeutic impact of melatonin on the fibrotic phenotype of HBdSMCs. Cells were co-treated for 72 hours with various dosages of melatonin (1, 10, 20, 50 uM) and TGF-β1 (10 ng/mL). Finally, we determined the optimal TGF-β1 concentration (10 ng/mL) and melatonin concentration (10 ng/mL) and used these experimental conditions as the standard for subsequent experiments.

### 2.4 Cell viability analysis

In our in vitro experiments, melatonin was solubilized in DMSO. To assess potential cytotoxicity to human bladder smooth muscle cells, we subjected the cells to varying concentrations of melatonin—1uM, 10uM, 20uM, and 50uM. Cell Counting-kit 8 (Vazyme, Nanjing, China) assessed TGF-β1 cells 72 hours post-stimulation.

### 2.5 Western blot analysis

HBdSMCs and rat bladder tissues were lysed with cell lysate (Beyotime, Jiangsu, China). Subsequently, the protein concentration was assessed and quantified using BCA (TaKaRa, Japanese). Immunoblotting was conducted on cell and tissue extracts (30 μg total protein/well) employing 10% SDS-PAGE (EpiZyme; PG113) and specific antibodies to reveal the levels of expressed proteins.

### 2.6 Reverse transcription PCR (R T–PCR) assay

According to the product instructions, total RNA of HBdSMCs treated with TGF-β1 and melatonin was extracted using the rapid RNA purification kit (Shanghai Yishan Biotech, Shanghai, China). Microultraviolet spectrophotometry was then used to determine the RNA’s concentration and purity. Following quantification, HiScript III RT SuperMix was used to create cDNA from total RNA. For qPCR, General Purpose High Sensitivity Dye-based Quantitative PCR Assay Kit (Q711, Vazyme) was employed to measure appropriate TGF-β1 and melatonin concentrations, as well as to evaluate cell viability. GADPH as the endogenous: Forward TTCACCACCATGGAGAAGGC, and reverse CTCGTGGTTCACACCCATCA; Collagen III: Forward TTCCTGGGAGAAATGGCGAC, and reverse GGCCACCAGTTGGACATGAT; α-SMA: Forward TGTGCTATGTCGCTCTGGAC and reverse AAGCGTTCATTCCCGATGGT

### 2.7 Histological analysis

Bladder tissue, fixed in 10% formalin, underwent paraffin wax embedding and subsequent cutting into continuous 5μm thick sections. To evaluate rat bladder fibrosis, sections were stained using HE and Masson trichromatic staining following standard protocols post-dewaxing and washing. The primary antibody was incubated on treated sections at 4°C for the immunohistochemical (IHC) staining procedure, which was followed by washing, the application of the secondary antibody coupled with HRP, and a final incubation at room temperature for an additional hour. Phospho-Smad2 (dilution 1:50, cat. AF3362, Affinity), Phospho-Smad3 (dilution 1:50, cat. AF3449, Affinity), N-cadherin (dilution 1:50, cat. 13116, CST), Vimentin (dilution 1:2500, cat. 60330, Proteintech), Collagen Type III (dilution 1:500, cat. 22734, Proteintech), E-cadherin (dilution 1:400, cat. 3195, CST), α-Smooth Muscle Actin (dilution 1:1500, cat. 14395, Proteintech), TGF-β1 (dilution 1:200, cat. 21898, Proteintech), CCN1 (dilution 1:100, cat. AF3362, Affinity).

### 2.8 Statistical analysis

Results were presented as mean ± standard error of the mean and analyzed with GraphPad Prism 9.0. Intergroup distinctions were evaluated through the unpaired t-test, one-way analysis of variance, and post hoc pairwise comparisons using Tukey’s test. P values below 0.05 were deemed statistically significant.

## 3. Results

### 3.1 Melatonin inhibits the TGF-β1-induced fibrotic phenotype of human bladder smooth muscle epithelial cells

To establish the TGF-β1-induced fibrosis model in HBdSMCs, cell cultures were conducted with varying concentrations of TGF-β1 (5, 10, and 20 ng/mL) for different durations (48h, 72h). Hence, Collagen III and α-SMA were employed as reference markers for fibrosis, and their expression was scrutinized using real-time PCR and Immunoblotting analysis. The findings revealed that when TGF-β1 concentration was 10ng/mL for 72 hours, the increase of Collagen Ⅲ was the most obvious. However, when TGF-β1 concentration was 20ng/mL for 72 hours, the increase of α-SMA was most obvious (Fig 1A). Because fibrosis is dominated by increased collagen fibrin, we used a TGF-β1 concentration of 10ng/mL and a culture time of 72h as the best reference conditions for subsequent fibrosis models. In this experiment, melatonin is dissolved in DMSO, and certain concentrations of DMSO have a toxic effect on cells. To investigate whether melatonin is cytotoxic to human bladder smooth muscle cells, we treated human bladder smooth muscle cells with melatonin (1uM, 10uM, 20uM, 50uM). The CCK8 assay showed that melatonin had no significant cytotoxic effect on human bladder smooth muscle cells at the above concentrations (S1B Fig). Following the identification of the optimal concentration (10 ng/mL) and duration (72h) of TGF-β1, we investigated the therapeutic impact of melatonin on the fibrotic phenotype of HBdSMCs. HBdSMCs induced by TGF-β1 (10 ng/mL, 72h) were subjected to melatonin treatment at concentrations of 1, 10, 20, and 50uM. PCR and immunoblot analysis demonstrated that, at a concentration of 50μmol/L melatonin, Collagen III and α-SMA exhibited the most substantial decrease (Fig 1B). To corroborate the impact of melatonin on TGF-β1-induced fibrosis, we specifically examined the outcome of melatonin at a dose of 50μmol/L (Fig 1C).

**Fig 1.**
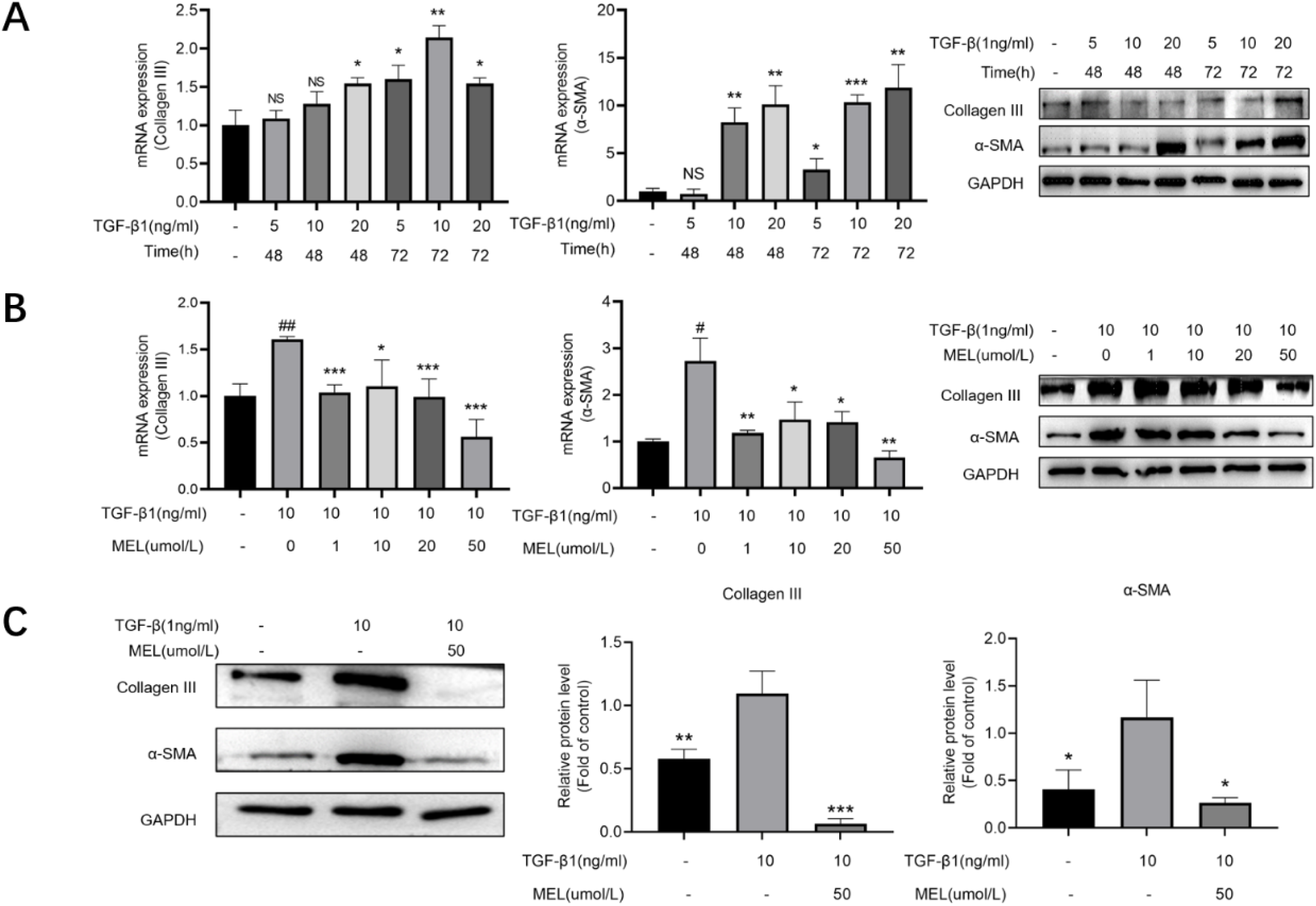
Effects of TGF-β1 on the expression of indicators fibrosis in HBdSMCs, and the effects of MEL on the fiber indicators of HBdSMCs induced by TGF-β1. (A) HBdSMC were inoculated in six-well plates with different levels of (5, 10 and 20 ng/mL) TGF-β1 and incubated for 48 h and 72 h. Cells were processed immediately after reaching the specified time, and the expressions of type III collagen and α-SMA proteins were detected by PCR and WB. (B) In the previous experiment, we determined the optimal TGF-β1 concentration of 10 ng/mL and the optimal action time of 72 h. Then HBdSMCs within 10 generations were inoculated into six-well plates, and different concentrations of melatonin were added (1uM, 10uM, 20uM, 50uM). total RNA and protein were extracted after 72h. The protein expression levels of type III collagen and α-SMA were detected by PCR and WB. Compared with the control group. (C) By the previous step of the experiments, we took the melatonin concentration (50uM) as the optimal treatment concentration. For subsequent experiments, we used TGF-β1 (10ng/mL, 72h) and melatonin concentration (50uM) as the basis, and on this basis, we detected the protein expression levels of type III collagen and α-SMA using WB. #P<0.05, ##P<0.01, *P<0.05, **P<0.01, ** *P<0.001 vs Control group

### 3.2 Melatonin regulates the TGF-β1/Smad pathway and EMT

To elucidate the mechanism by which melatonin inhibits TGF-β1-induced fibrosis in HBdSMCs, we conducted a comprehensive investigation into the alterations in the TGF-β1/Smad signaling pathway and EMT. The ideal TGF-β1 induction conditions (10 ng/mL, 72 hours) and melatonin treatment concentration (50 mol/L) were used to validate subsequent research. Evaluations were made of the relevant proteins’ nexpression. P-Smad2 and P-Smad3 expression in the group of models was higher than in untreated cells, and melatonin (50 mol/L) significantly decreased the expression of these proteins (Fig 2A). Smad2/3 expression remained unchanged under all treatment factors. When compared to untreated cells, the expression of Vimentin and N-cadherin rose in the model set, but E-cadherin expression dropped (Fig 2B). Melatonin (50 mol/L) significantly reversed the expression of these proteins. According to our results, melatonin substantially reduces the TGF-β1-induced fibrotic in HBdSMCs by modifying the TGF-β1/Smad pathway and EMT.

**Fig 2.**
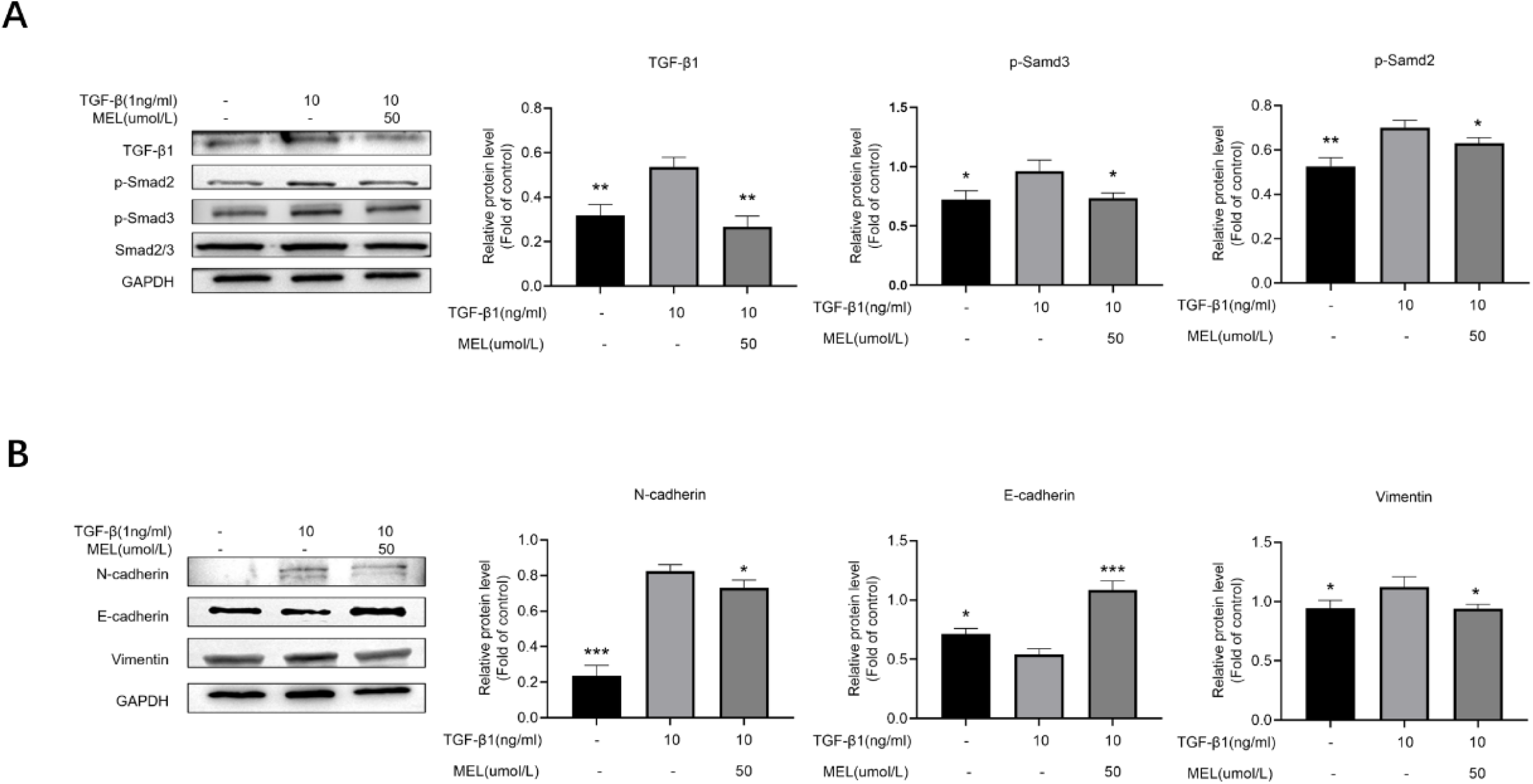
Effect of melatonin (50uM) on TGF-β1 (10ng/mL, 72h) mediated TGF-β1/Smad pathway and EMT in HBdSMCs. (A) HBdSMCs up to 10 generations were inoculated in six-well plates and experiments were performed under the above conditions. proteins were extracted after 72h and the expressions of TGF-β1, p-Smad2, p-Smad3 and Smad2/3 were detected by WB. (B) HBdSMCs within 10 generations were inoculated in six-well plates and experiments were performed under the above conditions. proteins were extracted after 72h and the expression levels of E-cadherin, N-cadherin and Vimentin were detected by WB. *P<0.05, **P<0.01, ***P<0.001 vs TGF-β1 group.

### 3.3 LPS induced bladder fibrosis in rats

A dual infusion of PS and LPS was employed to induce bladder tissue fibrosis in rats (Fig 3A, B). To assess whether this led to bladder wall hypertrophy and fibrogenesis, we conducted histological and morphometric analyses using HE and Masson trichrome staining. Following 6 weeks of perfusion, examination under HE staining revealed notable atypical hyperplasia of the bladder epithelium and hyperplasia of connective tissue in the bladder submucosa compared to the sham group (Fig 3C). Masson’s trichrome staining results indicated that the LPS group exhibited a greater degree of bladder smooth muscle hypertrophy and conspicuous collagen deposition compared to the sham group (Fig 3C and Fig S2A), in consistent with the data from Chae-Min Ryu(13).

**Fig 3.**
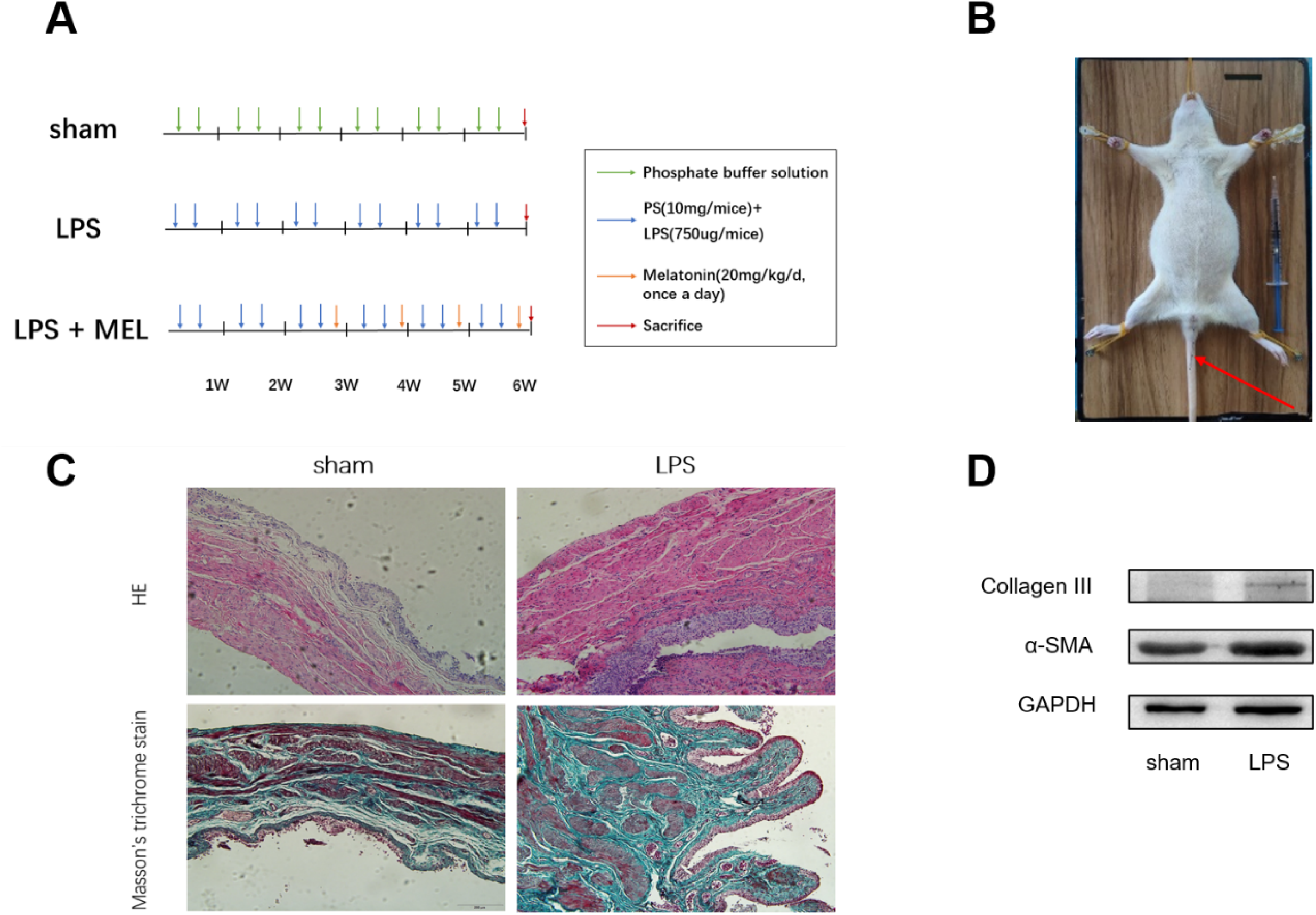
Histologic and protein changes in the PS/LPS-induced bladder fibrosis model in rats. (A) Schematic diagram of the experimental flow design of rat bladder fibrosis. (B) SD rat model (1 mm epidural catheter at red arrow). (C) HE staining (obvious change in bladder thickness) and Masson trichrome staining (collagen fibers in blue) of bladder tissues of rats with different treatments, 100×. (D) Immunoblot analysis for expression of collagen III and α-SMA in PS/LPS-induced rat bladder tissue.

This outcome was corroborated through immunoblotting (Fig 3D and S2B Fig). Thus, these data suggest that double instillation of LPS can effectively induce fibrosis in rat bladder tissue.

### 3.4 Melatonin reduces PS/ LPS-induced bladder fibrosis in rats

We went on to assess the effect of melatonin on the histological changes induced by LPS double perfusion. First, in the current model, HE in the LPS instillation group showed marked atypical epithelial hyperplasia of the bladder and connective tissue hyperplasia of the bladder submucosa, which was consistent with our previous results (Fig 4A). HE staining following melatonin treatment revealed a significant reduction in atypical epithelial hyperplasia of the bladder and connective tissue hyperplasia in the bladder submucosa compared to the LPS group, suggesting that melatonin mitigated bladder fibrosis (Fig 4A). Immunoblotting and immunohistochemistry were used to measure the expression of associated proteins. Collagen III and –SMA expression levels significantly outpaced the levels of the sham group following LPS/PS treatment (Fig 4B, C). After melatonin administration, the level of these proteins’ expression was noticeably reduced (Fig 4B, C). These findings suggest that melatonin effectively mitigates LPS/ PS-induced bladder fibrosis in rats.

**Fig 4.**
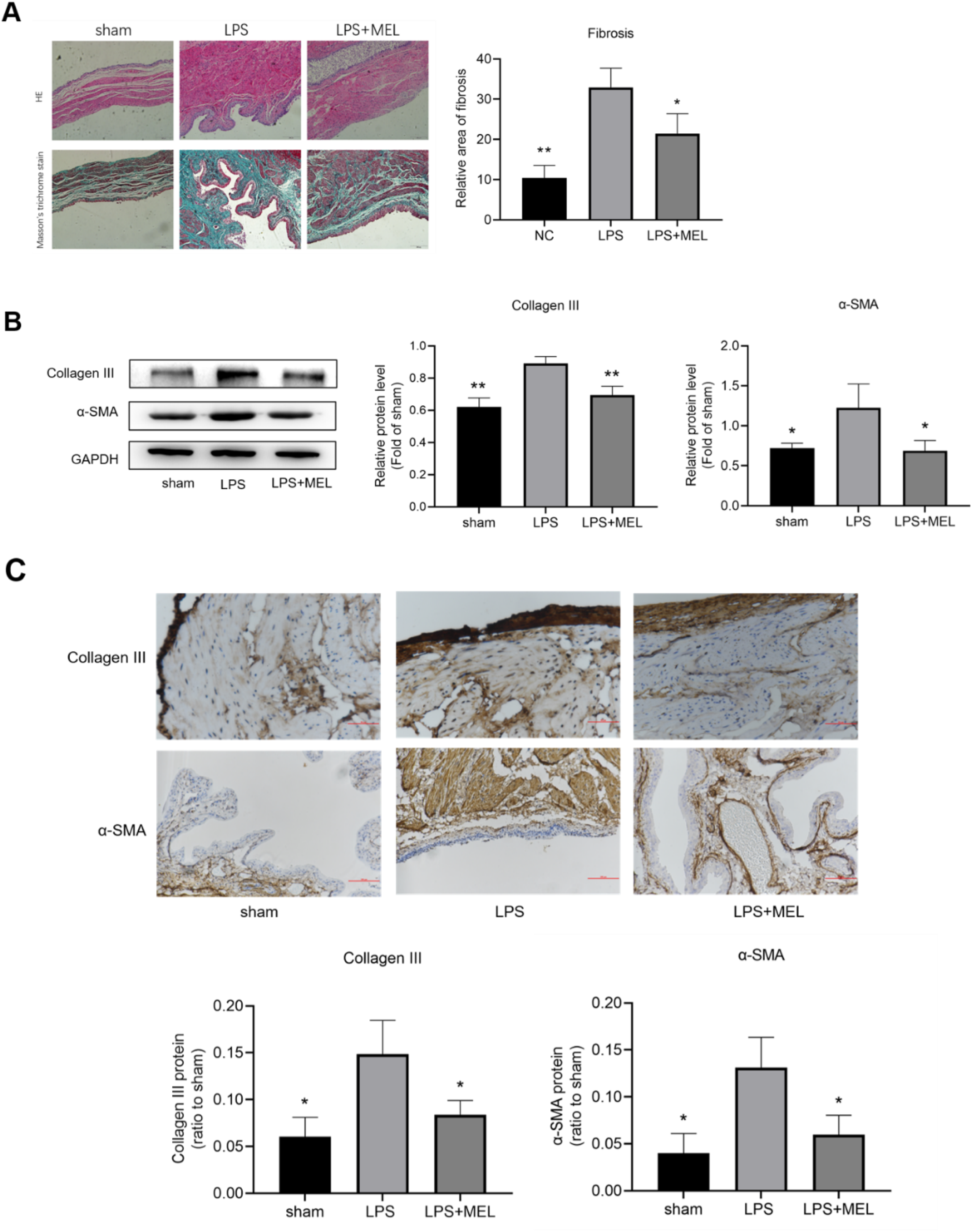
Therapeutic effect of melatonin on PS/ LPS-induced bladder fibrosis in rats. (A) Rat bladder tissue under different experimental conditions. Representative results of H&E and Masson trichrome staining, 100X. (B) Expression of collagen Ⅲ and α-SMA in rat bladder tissues under different experimental conditions using WB. (C) Representative immunohistochemical results of collagen III and α-SMA in rat bladder tissues under different experimental conditions. Collagen III, 400X; α-SMA, 200X. *P<0.05, **P<0.01, ** *P<0.001 vs LPS group

### 3.5 Melatonin regulates the TGF-β1/Smad signaling pathway and EMT in PS/ LPS-induced fibrotic bladder of rats

To further investigate the impact of melatonin on the TGF-β1/Smad signaling pathway and EMT in rat bladder tissues, we measured the expressions of related proteins using immunohistochemistry and immunoblot analysis. TGF-β1, p-Smad2 and p-Smad3 expression intensity was considerably increased after PS/LPS stimulation compared to the sham-operated group, whereas it was significantly lower after melatonin therapy (Fig 5A). Smad2/3 expression was constant across all groups. Similar findings from an immunohistochemical study were obtained (Fig 5B). These results indicate that the rat bladder fibrosis model’s degree of fibrosis can be attenuated by melatonin through decreasing the expression of the TGF-β1/Smad signaling pathway. To determine the status of EMT, we identified pertinent proteins in bladder tissues using immunohistochemistry staining and immunoblot analysis. The findings demonstrated that whereas the expression of E-cadherin reduced in the LPS group relative to the sham group, the level of N-cadherin and vimentin increased. When compared to the LPS group, these proteins were reversed in the bladder tissues of the melatonin-treated rats (Fig 5C, D). The above findings show that Melatonin can significantly slow down EMT and ameliorate the fibrosis process in the PS/LPS fibrosis model, which highlights the importance of this mechanism.

**Fig 5.**
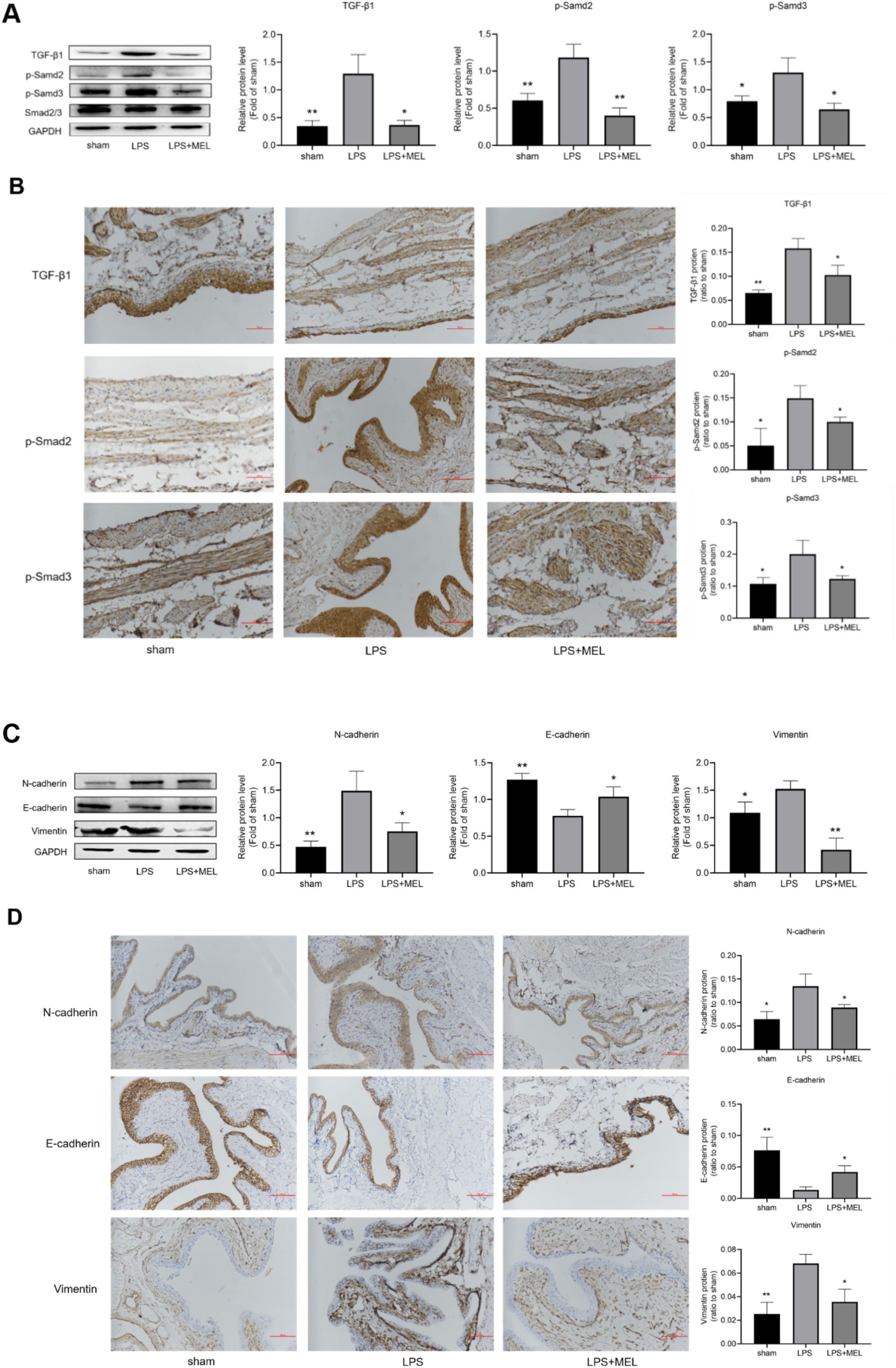
Effects of melatonin on TGF-β1/Smad pathway and EMT in PS/LPS-induced bladder fibrosis model in rats. (A, B) Representative results of WB and immunohistochemistry of TGF-β1, p-Smad2, p-Smad3 and Smad2/3 in rat bladder tissues under different experimental conditions. (C, D) Representative WB and immunohistochemical results of E-cadherin, N-cadherin and Vimentin in rat bladder tissues under different experimental conditions. All protein indexes, 200X *P<0.05, **P<0.01, ** *P<0.001 vs LPS group.

### 3.6 Quantitative proteomic analysis of cells with increased fibrosis phenotype and melatonin treatment

To decipher the proteins and biological pathways implicated in the anti-fibrotic effects of melatonin, we conducted proteomic analysis using human bladder smooth muscle cells from the control group, TGF-β1 group, and TGF-β1+ MEL group (n = 3 for each group, Fig 6A). A total of 6100 non-redundant proteins were quantified by LC-MS/ MS-based proteomics experiments (Table S1). All the groups exhibited good biological reproducibility, as evidenced by the Pearson correlation coefficient, which measured the correlation between any two samples in each group, being greater than 0.94 (S3A Fig). When comparing the TGF-β1 simulation group with the control group, the findings showed substantial changes in 832 proteins. 212 of these proteins showed up-regulation, whereas 620 showed down-regulation (Fig 6B). Between the MEL treatment group and the TGF-β1 model, an aggregate of 1564 proteins were substantially different, of which 996 proteins were elevated and 568 proteins were down-regulated. Subsequently, we identified the top 9 proteins exhibiting the most substantial increase in the TGF-β1 model. Similarly, we identified the top 9 proteins with the most decrease in expression in the MEL-treated group compared to the TGF-β1 model (Fig 6C). Combining the two sets of data (Fig 6D), we found that CCN1 was most likely the target protein of melatonin treatment. Western blot analysis was employed to authenticate CCN1 expression in diverse experimental cohorts. Remarkably, CCN1 expression significantly heightened in both the TGF-β1 model and LPS groups. In contrast, CCN1 expression diminished in melatonin-treated human bladder smooth muscle cells and rat bladder tissue. Immunohistochemical analysis corroborated the Western blot findings, indicating an increase in CCN1 expression in the LPS group, followed by a reduction post-melatonin treatment (Fig. 6D). These outcomes support CCN1 as a pivotal protein in bladder fibrosis progression, with melatonin demonstrating efficacy in reducing CCN1 expression and ameliorating fibrotic severity.

**Fig 6.**
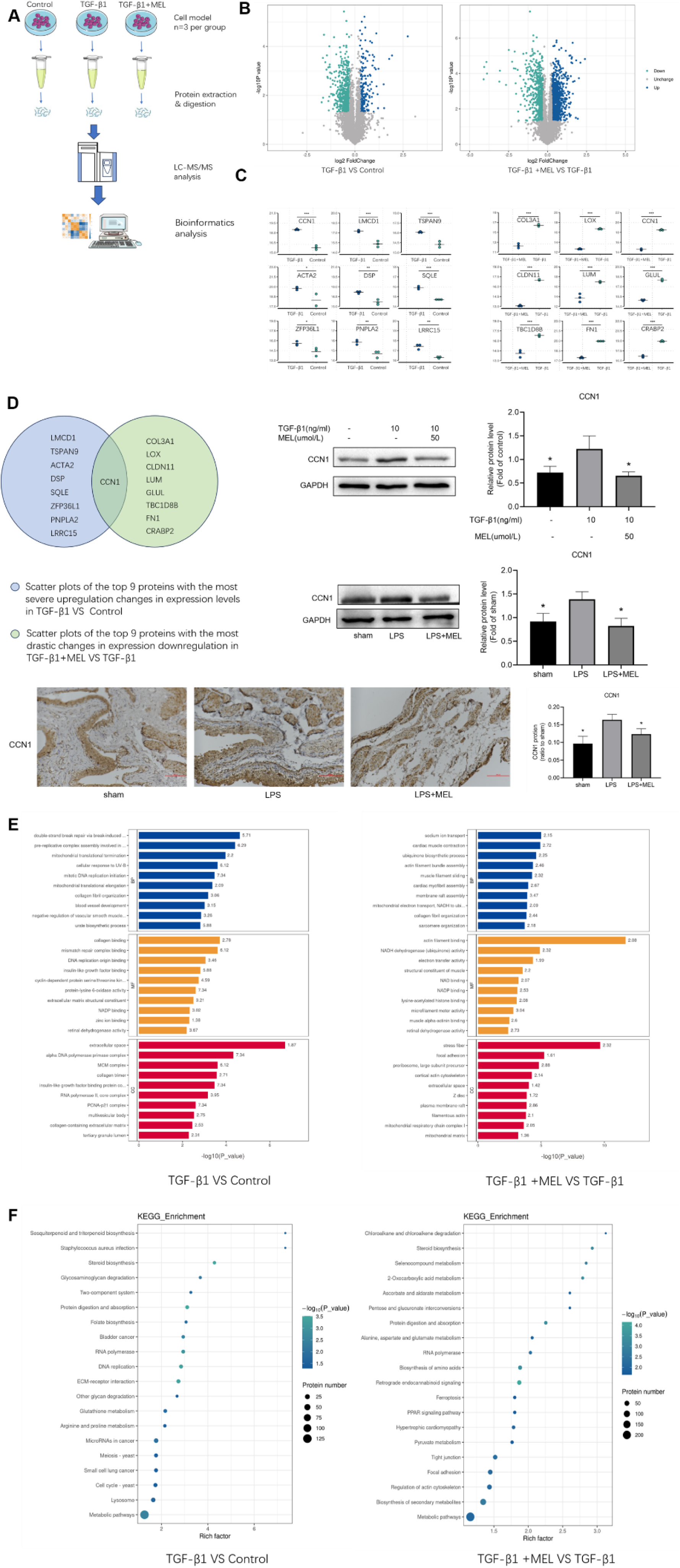
Proteomic analysis of control, TGF-β1 model, and melatonin-treated HBdSMCs. (A) Workflow diagram of label-free quantitative proteomics experiments. Total proteins were extracted from different groups of cells (control group, TGF-β1 model group and melatonin group, n = 3 each).and digested with trypsin. Equal amounts of separated peptides were analyzed by LC-MS/MS. The specific protein database was used for bioinformatics analysis of differentially expressed proteins. (B) The results of differential protein screening were displayed in the form of volcano plot. Dots in the figure, blue denotes up-regulated proteins, cyan denotes down-regulated proteins, and gray-black denotes no difference. The abscissa is the fold change (base 2 log transformation). The ordinate is the P-value value (base 10 log transformed). (C) The top 9 most up-regulated proteins between TGF-β1 model and control group and the top 9 most down-regulated proteins between MEL treatment group and TGF-β1 model. (D) CCN1 may be a targeted protein for melatonin treatment, and the expression of CCN1 in each group was detected by WB under different experimental conditions, and the expression of CCN1 in rat bladder tissue was verified by immunohistochemistry. (E) Bar graphs show GO enrichment analysis of differential proteins. (F) The top 20 enriched pathways in KEGG enrichment analysis results. *P< 0.05, **P<0.01, ** *P<0.001 vs TGF-β1 group

Substantial enrichment analysis of Gene Ontology (GO) functional annotations revealed that the proteins exhibiting differential expression before and after TGF-β1 treatment were primarily linked to the extracellular space in cellular composition. In contrast, the differential proteins before and after melatonin treatment were predominantly associated with actin filament binding and stress fiber-related molecular functions in cellular composition (Fig 6E). Moreover, through an analysis of KEGG pathway enrichment for differential proteins among various treatment groups, we observed that these proteins were primarily concentrated in metabolic pathways (Fig 6F). Hence, these results lead us to speculate that addressing the structural aspects of the extracellular space linked to metabolic pathways, actin filament binding, and stress fibers could be crucial focal points for fibrosis treatment.

## 4. Discussion

Bladder fibrosis is a common pathological process related to many urinary system diseases. In addition to BPH-related BOO, many other diseases have been associated with bladder fibrotic changes, such as aging stroke, Parkinson’s disease, multiple sclerosis, and diabetes (14). Cystic fibrosis is a challenging issue that hasn’t been solved yet since bladder function is weakened and difficult to recover from after long-term chronic harm to the bladder (15).The rat bladder fibrosis model in this study referred to the PS/LPS double perfusion method established by Chae-Min Ryu et al., and two weeks of perfusion time was added on this basis, because the histological appearance of this model was more consistent with Hunner type IC/BPS related bladder lesions and more meaningful for clinical treatment.

Bladder fibrosis manifests as the proliferation of BSMCs and the accumulation of ECM, resulting in compromised bladder fluid storage and emptying functionality(16, 17). Fibrotic BSMCs can transition from a contractile, nonproliferative phenotype to a synthetic one, altering the interaction between BSMCs and ECM and resulting in the contractile dysfunction of BSMCs (18). Remodeling of the ECM and increased expression of TGF-β1 are considered the mechanisms underlying bladder fibrosis (19).

A three-dimensional macromolecular network made up of collagen, fibronectin, and various other glycoproteins makes up the bladder ECM. A complex network is created when matrix elements interact with one another, and cells cling to adhesion receptors. The regular maintenance of homeostasis depends on the regulation of vital cellular processes like migration, growth, survival, and differentiation by cell surface receptors, which translate signals from the external matrix into cells. Under both normal and pathological circumstances, a number of different matrix-degrading proteins constantly remodel the ECM, which is a fluid architectural system(20). Collagen types I and III, in fibrous form, serve as the principal scaffold matrix proteins in the ECM, contributing to structure, tensile strength, and bladder compliance in fibrotic diseases (21). A substantial rise in type III collagen may arise from enhanced synthesis, with excessive collagen deposition marking the concluding phase of the fibrotic process. The Collagen fibers in the parietal layer of the bladder are mainly composed of Collagen Ⅰ, Collagen Ⅲ and Collagen Ⅳ. Among them, the content of Collagen Ⅲ is the factor that determines the bladder compliance. The more collagen III is deposited, the lower bladder elasticity leading to bladder dysfunction(22, 23). Through integrin-cytoskeleton and other mechanisms, highly elastic and low viscous extracellular matrix promotes the production of –SMA during the advancement of fibrosis (24, 25). Results indicated that these protein in HBdSMCs treated with TGF-β1 and rats treated with LPS exhibited higher levels compared to the control group.

There are three different TGF-βisoforms (TGF-β1, TGF-β2, and TGF-β3) found in animals, according to research. All three TGF-β isoforms interact with the TGFR2 receptor, which then attracts and activating the TGFR1 receptor(26). Smad2 and Smad3 are activated by TGFβ-R1, which causes them to separate from the type I receptor and form a heterotrimeric complex with Smad4 that translocates into the nucleus. Smad, in collaboration with general transcriptional factors, other transcription factors, or auxiliary proteins, governs the transcription of target genes (9, 27, 28).

EMT is an intricate process governed by an extensive interactome, comprising protein-protein and gene interactions triggered and regulated in response to extracellular signals. At the forefront of these interactions is TGF-β1. TGF-β1 exhibited a promoting effect on SNAIL1. ZEB1 is a downstream gene of SNAIL1, and SNAIL1 also enhances ZEB1. SNAIL1 and ZEB1 inhibit E-cadherin and promote the expression of N-cadherin and Vimentin(29). These findings are consistent with our experimental results.

Melatonin possesses sleep-inducing qualities and controls seasonal and circadian rhythms. Being a multifunctional molecule, it also possesses other qualities like anti-inflammatory and immunomodulatory ones (30, 31). The fibrotic response primarily encompasses four stages: initial organ injury, activation of effector cells, formation, and dynamic deposition of the extracellular matrix (ECM). Melatonin modulates each of these stages, and additionally, it has the capacity to diminish fibrosis levels in various organs(12). The therapeutic efficacy of melatonin against LPS-induced bladder fibrosis in rats in this experiment is substantiated by the mitigation of histopathological observations. The TGF-β1/Smad pathway was assessed through immunohistochemistry and Western blotting. We noted elevated expression of TGF-β1, p-Smad2, and p-Smad3 in the double perfusion group compared to the sham operation group. Nevertheless, melatonin alleviates these effects. Similar outcomes were noted in the in vitro cellular model. These data imply that the administration of melatonin leads to the downregulation of the TGF-β1/Smad pathway. Simultaneously, we adopted the same approach to assess the expression of EMT. We noted an increase in the expression of N-cadherin and vimentin, coupled with a decrease in E-cadherin in the LPS double perfusion group compared to the sham operation group. Simultaneously, these trends improved with melatonin treatment. These results indicate that melatonin administration has the capability to alleviate EMT. Consequently, we infer that the preventive impact of melatonin on bladder fibrosis might be associated with the inhibition of TGF-β1/Smad pathway and EMT.

We conducted proteomic analysis utilizing HBdSMCs from the control, TGF-β1 model, and melatonin-treated groups. We identified CCN1 as a probable target protein for melatonin in the treatment of fibrosis. CCN1 is a multifunctional protein, playing a pivotal role in diverse physiological processes, including embryonic development, aging, vascular maintenance, and tissue damage repair (32, 33). Additionally, CCN1 has been linked to a few clinical conditions, such as cancer, atherosclerosis, arthritis, and fibrosis (34–37). CCN1 was activated by lots of growth factors including TGF-β1, FGF2, GH, etc(38). In contrast to normal human lung tissue, Kurundkar et al. found a significant increase of CCN1 expression in the lung tissues of individuals with idiopathic pulmonary fibrosis. After that, they demonstrated through a number of experimental investigations that CCN1 causes lung fibrosis by boosting the expression of the TGF-1/SMAD3 pathway (39). Kulkarni et al. found that CCN1 could mediate pro-fibrotic effects by augmenting TGF-β1 signaling (40). ZHI-QIANG LI et al. also observed a continuous increase in CCN1 expression in liver fibrosis, suggesting a potential association with the progression of liver fibrosis (41). We have found through experimental studies that exogenous TGF-β1 activated CCN1, potentially contributing to bladder fibrosis via the TGF-β1/SMAD pathway. Nevertheless, certain studies have indicated that CCN1 can exert an anti-fibrotic role by fostering fibroblast senescence and apoptosis(42). Consequently, additional research is required to elucidate the precise relationship between CCN1 and fibrosis. Furthermore, squalene cyclooxygenase (SQLE), identified as a ferroptosis gene, exhibited a significant reduction following melatonin treatment. Consequently, SQLE may represent another target protein for melatonin in the treatment of fibrosis. SQLE is a crucial enzyme in cholesterol biosynthesis, and no pertinent reports on SQLE in fibrosis have been documented. Nevertheless, there are reports indicating that SQLE propels the proliferation of colon cancer cells, fosters intestinal dysbiosis, and expedites colorectal cancer(43). More intriguingly, studies have also uncovered that the diminution of squalene epoxide hydrolase can expedite the progression and metastasis of colorectal cancer (44). In a recent study, Zhirui Zhang et al. identified a significant upregulation of SQLE expression in hepatocellular carcinoma (HCC), which is associated with an adverse clinical prognosis. SQLE promotes HCC growth, epithelial-mesenchymal transition, and metastasis by activating TGF-β/SMAD signaling (45). We found by immunoblotting that SQLE exhibited a substantial increase in the TGF-β1 group and the LPS group, whereas its expression significantly decreased following MEL treatment (S3B Fig). Hence, SQLE represents another pivotal protein that will be a focal point in our future investigations.

Our study yielded the following primary findings: firstly, we established a rat model of chronic inflammation-induced bladder fibrosis through continuous intravesical instillation of PS and LPS over a 6-week period; Secondly, LPS activation induces bladder injury via the TGF-β1/Smad and EMT pathways, which play pivotal roles in the pathogenesis of bladder fibrosis. Thirdly, melatonin modulation of the associated TGF-β1/Smad pathway and EMT proved effective in alleviating chronic inflammation-induced bladder fibrosis in rats. However, this study is subject to certain limitations. Firstly, the outcomes of this study are centered on fibrosis induced by chronic inflammation. The therapeutic efficacy of melatonin was not assessed in alternative bladder fibrosis models, including neurogenic and outlet obstruction models. Secondly, given the intricate pathogenesis in patients with bladder fibrosis, these investigations have been confined to in vitro and animal studies. There is a considerable distance to cover before these treatments translate into clinical health benefits, and the translational relevance of experimental findings to humans remains constrained. Lastly, the subsequent functional verification and mechanistic investigation of CCN1 have yet to be undertaken, and relevant efforts are currently underway in our laboratory.

The pathophysiological mechanism underlying bladder fibrosis remains incompletely elucidated, and the therapeutic outcomes are suboptimal. The pathophysiological mechanisms of bladder fibrosis are not fully understood, and current treatments are ineffective. In this study, melatonin-based treatment demonstrated a mitigating effect on fibrosis in a rat model of LPS-induced fibrosis. Given the limited therapeutic options for chronic inflammation and fibrosis of the bladder, melatonin might represent a novel therapeutic approach, with CCN1 potentially serving as a key therapeutic target.

## 5. Conclusions

Melatonin is a promising new treatment in chronic cystitis and bladder fibrosis, and CCN1 serves as a key therapeutic target.

## Acknowledgements

Not applicable.

## Supporting information

**S1 Fig.**
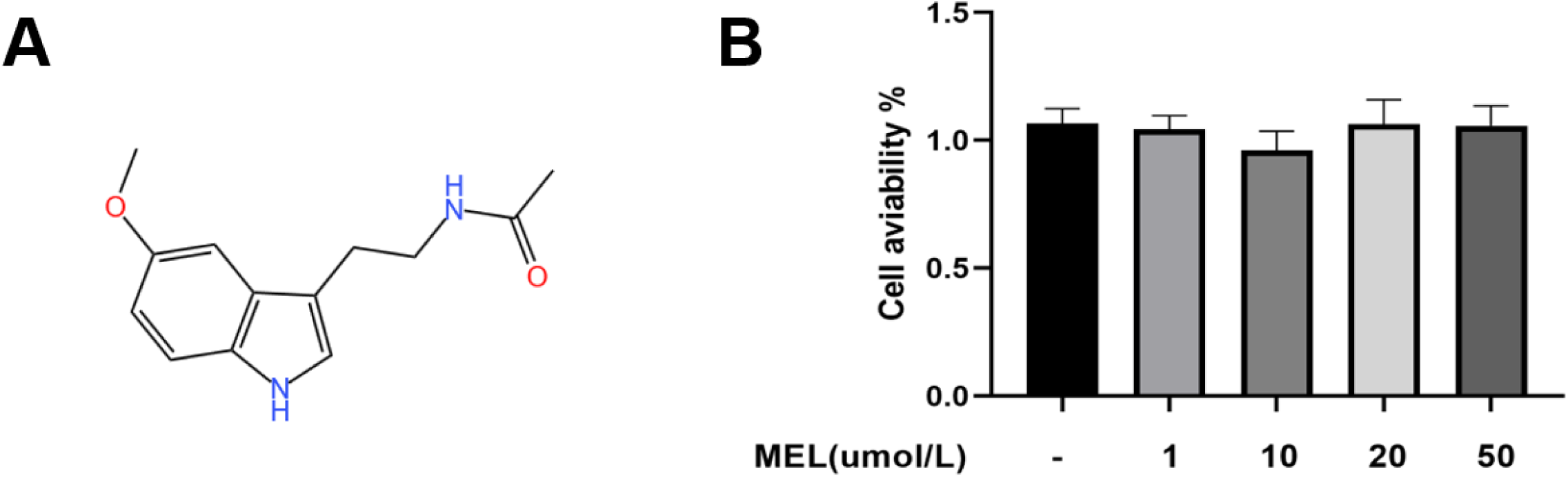
(A) Chemical structural formula of melatonin. (B) CCK-8 detection of the effect of different concentrations of melatonin on the activity of HBdSMCs.

**S2 Fig.**
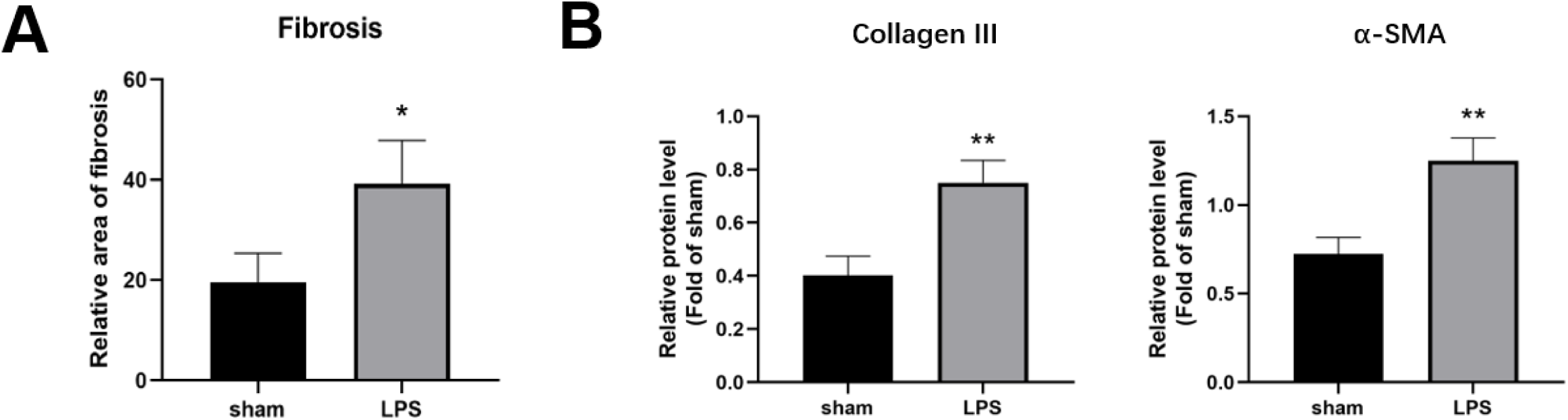
(A) Statistical analysis of data after quantification of Masson trichrome staining of rat bladder tissues from sham and double perfusion groups. (B) Statistical analysis of data after quantification of collagen III and α-SMA expression in rat bladder tissues by WB assay. *P<0.05, **P<0.01 vs sham group.

**S3 Fig.**
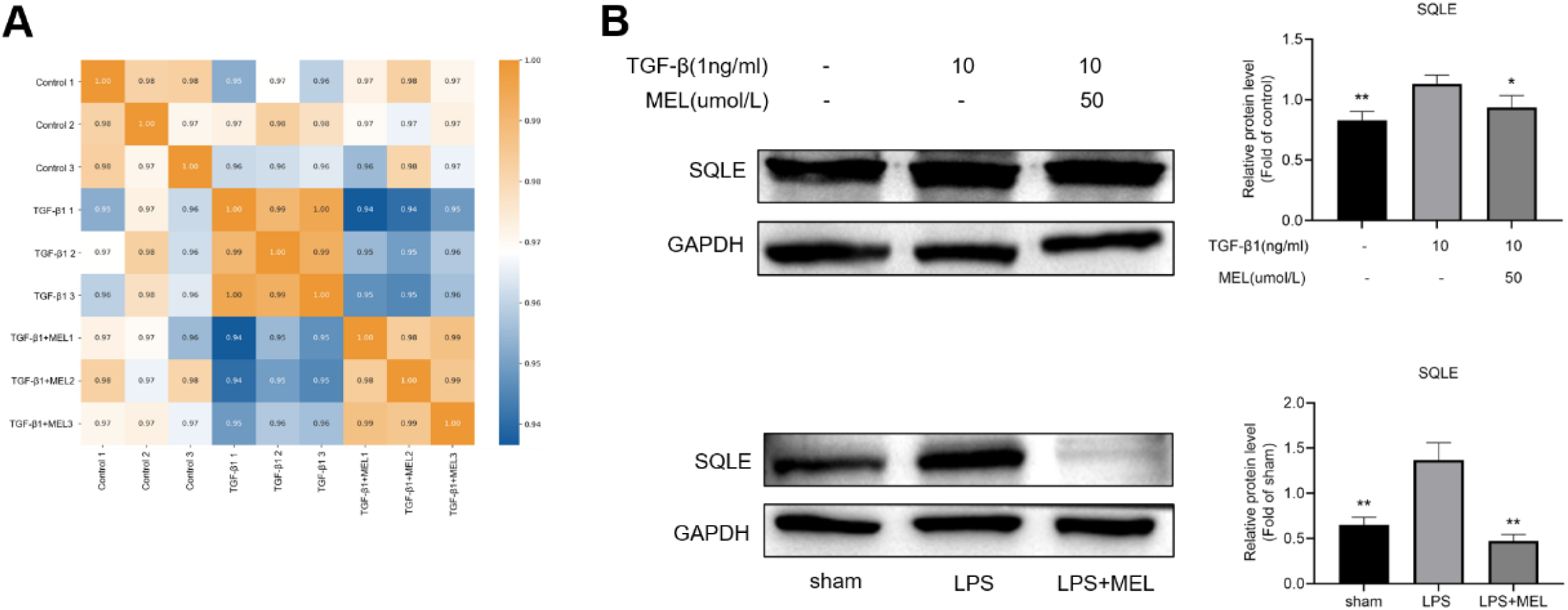
(A) Heat maps were plotted by calculating the Pearson correlation coefficient between all sample pairs. (B) WB detection of changes in SQLE protein expression in HBdSMCs and rat bladder tissues. *P<0.05, **P<0.01 vs model group

